# The intratumoral microbiome is negatively associated with malignancy in dermatofibrosarcoma protuberans

**DOI:** 10.1101/2025.06.02.657449

**Authors:** Xingxing Jian, Ziyang Liu, Jingtian Zhang, Yani Li, Lu Xie, Guihu Zhao, Bin Li, Jinchen Li, Cong Peng, Miaojian Wan, Xiang Chen

**Affiliations:** Bioinformatics Center, National Clinical Research Center for Geriatric Disorders, Department of Geriatrics, Xiangya Hospital, Central South University, Changsha, Hunan, China; Department of Dermatology, Xiangya Hospital, Central South University, Changsha, Hunan, China; Department of Dermatology, The Third Affiliated Hospital, Sun Yat-sen University, Guangzhou, Guangdong, China; Hunan Key Laboratory of Skin Cancer and Psoriasis, Hunan Engineering Research Center of Skin Health and Disease, Xiangya Hospital, Central South University, Changsha, Hunan, China; Furong Laboratory, Changsha, Hunan, China; National Clinical Research Center for Geriatric Disorders, Xiangya Hospital, Central South University, Changsha, Hunan, China

**Keywords:** Dermatofibrosarcoma protuberans (DFSP), Intratumoral microbes, Microbiome, Whole-genome sequencing (WGS), *Streptomyces* spp

## Abstract

In recent years, growing evidence demonstrated the critical roles of intratumoral microbes in tumors. However, the characteristics of intratumoral microbiome in dermatofibrosarcoma protuberans (DFSP) and their associations with tumor malignancy remains unexplored. Here, 106 whole-genome sequencing (WGS) data of the paired biopsy and blood samples from 53 DFSP patients were utilized, in which we extracted high-quality non-human reads to assign microbial taxonomic annotation by Kraken2 and PlusPF database. Meanwhile, significantly elevated non-human reads, microbial abundance, and bacterial load were detected in tumor compared to blood samples. To minimize the impact of contaminants and noise, we conducted a series of filtering procedures and treated blood samples as negative controls. Notably, we found that over 98% of intratumor microbes belonged to bacteria, and that the dominant taxa and alpha-diversity of intratumoral bacteria all exhibited significantly negative correlations with Ki67 expression in tumors. Specifically, the 84 key intratumoral species were identified to be associated with the reduced tumor malignancy, as evidenced by the diminished Ki67 expression, decreased tumor volume, and fewer *COL1A1-PDGFB* fusion events. These findings highlighted the tumor-suppressive potential of intratumoral microbiome, thereby providing a novel insight into DFSP pathogenesis and clinical management.

**IMPORTANCE:** To our knowledge, this is the first study to explore the intratumoral microbiome in DFSP. Microbial DNA signatures could be detected by whole-genome sequencing (WGS) and enabled a comprehensive view of the microbial landscape, thus we analyzed 106 WGS data of the paired biopsy and blood samples from 53 DFSP patients. Our analysis revealed that the dominant taxa and alpha-diversity of intratumoral microbiome all showed significantly negative correlations with Ki67 expression of tumor samples. This study indicated the tumor-suppressive role of intratumoral microbiome in DFSP and highlighted the potential clinical significance in understanding DFSP pathogenesis and guiding therapeutic management strategies.

## Background

Dermatofibrosarcoma protuberans (DFSP) is a cutaneous soft tissue sarcoma, with an incidence of approximately 6 cases per million individuals. ^[1]^ The pathogenesis of DFSP primarily involves the translocation between chromosomes 17 and 22, resulting in *COL1A1-PDGFB* fusion in approximately 87% of cases. ^[2]^ A history of trauma is also considered as a key factor in DFSP development, revealing a strong correlation between tumor growth and trauma associated DFSP. ^[3]^ Surgical resection remains the primary treatment modality for DFSP, aiming to achieve complete tumor excision while concurrently mitigating recurrence risk and preserving optimal functional and cosmetic outcomes. ^[4]^ And, a multidisciplinary approach may be considered in advanced cases, including surgery, immune checkpoint inhibitor, and radiotherapy. ^[5, 6]^ Despite the generally favorable prognosis and low metastatic potential in DFSP patients, it remains a critical priority to elucidate novel pathogenesis and mitigate local recurrence risk. ^[7]^

Thanks to advancements in high-throughput sequencing technology and bioinformatics tools, human microbiome has been extensively studied to enhance our understanding of microbial involvement in human health and disease. Actually, various regions of the human body, including blood and tumor tissues, harbor diverse microorganisms that serve as effective biomarkers. ^[8-10]^ In recent years, an increasing number of studies have highlighted the critical roles of intratumoral microbes in cancer pathogenesis, progression, and therapeutic response. ^[11]^ Therein, intratumoral microbes could influence tumor cells through three potential mechanisms: 1) immunomodulation via the immuno-oncology-microbiome (IOM) axis, 2) metabolic disturbance driven by microbial production of bioactive metabolites, and 3) disruption of cell adhesion by contact-dependent interactions. ^[12, 13]^ For example, *Bifidobacteria* was reported to enhance dendritic cell function and prime CD8^+^T cells, thereby promoting antitumor immunity and improving anti-PD-L1 efficacy in melanoma. ^[14]^ Secondary bile acids and short-chain fatty acids produced by gut microbiota as ligands were able to regulate metabolic reprogramming. ^[15]^ *Fusobacterium nucleatum* was revealed to colonize tumor tissue by attaching lectin Fap2 to tumor-displayed Gal-GalNAc, thereby promoting tumor growth and metastatic progression in breast cancer. ^[16]^ However, to date, no studies have explored the characteristics of intratumoral microbiome in DFSP.

Whole-genome sequencing (WGS) could detect DNA signatures from microbial genomes in addition to the human genome, enabling a comprehensive view of the microbial landscape in tumor. ^[10]^ In this study, based on the WGS data of paired biopsy and blood samples from DFSP patients, ^[17]^ those high-quality non-human reads were extracted and utilized to annotate microbial taxonomy. We observed that the predominant intratumoral taxa across the phylum, family, and genus levels, and alpha-diversities of intratumoral microbiome all exhibited negative correlations with Ki67 expression in DFSP. Several key intratumoral species, such as *Streptomyces* spp, were identified as representative. This present study initially investigated the role of intratumoral microbiome in inhibiting the tumor progression of DFSP, thereby providing an insight in DFSP management.

## MATERIALS AND METHODS

### Data source and participants

In this study, the 106 WGS data of the paired tumor and blood samples of 53 DFSP patients were obtained from a prior study. ^[17]^ Moreover, the available clinical information of these patients was also collected, including age, sex, duration of disease, Ki67 expression, tumor volume, trauma history, and *COL1A1-PDGFB* fusion events **(Table S1)**. Note that the tumor mutational burden (TMB), variant types, and SNV classes (T > G, T > C, T > A, C > T, C > G, C > A) yield in the previous study were also collected. ^[17]^ And, 43 tumor samples from these DFSP patients were subjected to fluorescence in situ hybridization (FISH) analysis by using the *COL1A1-PDGFB* fusion gene probe. ^[17]^

### Taxonomic annotation of intratumor and blood microbes

The WGS files of the paired tumor and blood samples were aligned with human genome (hg38) by BWA mem (v0.7.17-r1188), and then those non-human reads were extracted by Samtools (v1.9). To obtain high-quality non-human reads, BBDuk (v39.01) was used for reads filtering, and then Kneaddata (v0.12.0) embedding Trimmomatic (v0.33) and Bowtie2 (version 2.2.3) were used to trim adapter and remove human reads again. Subsequently, Fastp (v0.23.4) was used to assess the quality of non-human reads, and those samples with Q20 and Q30 above 0.95 and 0.85, respectively, were retained. Eventually, the high-quality non-human reads were applied for taxonomic annotation by Kraken2 (v2.1.3) and PlusPF database which includes human genome and microbial reference genomes from bacterial, viral, archaeal, fungal kingdoms. ^[18, 19]^ The workflow of taxonomic annotation above was summarized in **Fig. S1**.

### Filtering procedures of contaminants and noise

Referring to a prior study, ^[10]^ we proposed a series of filtering procedures to identify key intratumor species (**Fig. S1**). Firstly, according to a list of 93 bacterial genera as reagent contaminants, ^[20]^ microbial species from intratumor or blood samples were excluded as contaminants if they belong to these 93 contaminant genera and had a detection rate of greater than 25%. And, those remaining intratumor or blood species strongly correlated to at least one of these identified contaminants were also removed as contaminants (Spearman’s correlation: ρ>0.7, p<0.05). ^[21]^ Spearman’s correlation between two species was calculated based on the centred log-ratio-transformed microbial relative abundance using the function *clr* of R package compositions (v2.0-8). ^[22]^ Following this, we eliminated intratumoral species with a detection rate below 25% as noise, while those with detection rates above 25% were divided them into two categories: species detected exclusively in tumor samples and species detected in both tumor and blood samples. Therein, those detected in both tumor and blood samples need to show significantly higher detection rate in tumors by using Fisher’s test (p<0.05). ^[23]^ Finally, those species detected exclusively in tumor and those species with significantly higher detection rate in tumor were combined as the intratumoral species of DFSP.

### Diversity analysis of intratumoral microbiome

According to the abundance of intratumoral microbiome in DFSP, we separately calculated the alpha-diversities using the R packages vegan (version 2.6.8), including Shannon and Species Number (SpeciesNo). Moreover, we estimated the Bray-Curtis distances between tumor samples based on their abundance of intratumoral microbiome using the R packages ape (version 5.8), and then principal coordinates analysis (PCoA) was applied for visualization.

### WGCNA and gradient series analysis

On the basis of the abundance of intratumoral microbiome in DFSP patients, we carried out weighted gene co-expression network analysis (WGCNA) using the R packages WGCNA (version 1.73) and flashClust (version 1.1.2). The optimal soft threshold was determined using the function *pickSoftThreshold* to convert the correlation matrix into an adjacency matrix, with a soft threshold (softPower) chosen based on a scale-free topology fit. A topological overlap matrix (TOM) was then computed, and ‘genes’ with similar abundance patterns were grouped into modules using average linkage hierarchical clustering in the function *blockwiseModules*. Subsequently, all modules marked by distinct colors were used to correlate with clinical indicators by using Spearman’s correlation.

Additionally, we divided the tumor samples into four groups based on the quartiles of their tumor volume. To identify species which exhibited a decreasing trend with tumor volume, we then set the number of intratumoral microbial clusters to 20 and performed gradient series analysis using the R package Mfuzz (version 2.62.0).

### Statistical Analysis

Since the non-human reads were considered to be proportional to the total reads in each WGS sample, and in order to minimize bias in subsequent comparisons, the number of non-human reads in each sample was normalized by the sequencing depth of human genome, as following:

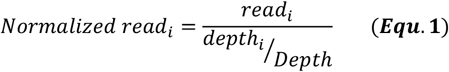

in which *read*_*i*_ and *Normalized read*_*i*_ separately stand for the non-human reads before and after normalization in the *i*-th sample; *depth*_*i*_ and *Depth* represent the sequencing depth of human genome in the *i*-th sample and the average depth, respectively. Here, the average depth of our 106 WGS files is 16x (range: 11x ∼ 23x).

All statistical analysis and visualization were carried out on the R platform (version 4.3.3). Significance was determined using Fisher’s test, t-test, and Wilcoxon test, while Spearman’s correlation was calculated using the R package psych (version 2.4.6.26).

## RESULTS

### Inferring microbial DNA signature in tumor and blood samples from DFSP patients

In this study, 106 whole-genome sequencing (WGS) data of paired biopsy and blood samples from 53 DFSP patients were obtained from a prior study. ^[17]^ To extract high-quality non-human reads and annotate microbial taxonomy in each WGS file, an improved bioinformatics workflow was proposed (**Methods**; **Fig. S1A**). ^[10]^ Initially, microbial sequences were detected extensively in tumors and blood of DFSP patients. We observed that the number of non-human reads increased with the sequencing depth of human genome, especially in tumor samples (**Fig. 1A**). This was speculated to result from the proportional generation of non-human reads to human reads during WGS. Thus, to reduce biases introduced by the distinct sequencing depths in WGS files, we normalized their non-human reads according to ***Equ. 1***. (**Fig. 1B**). Remarkably, we observed that the number of non-human reads in tumors was always higher than those in blood across the sequencing depths.

**Fig. 1.**
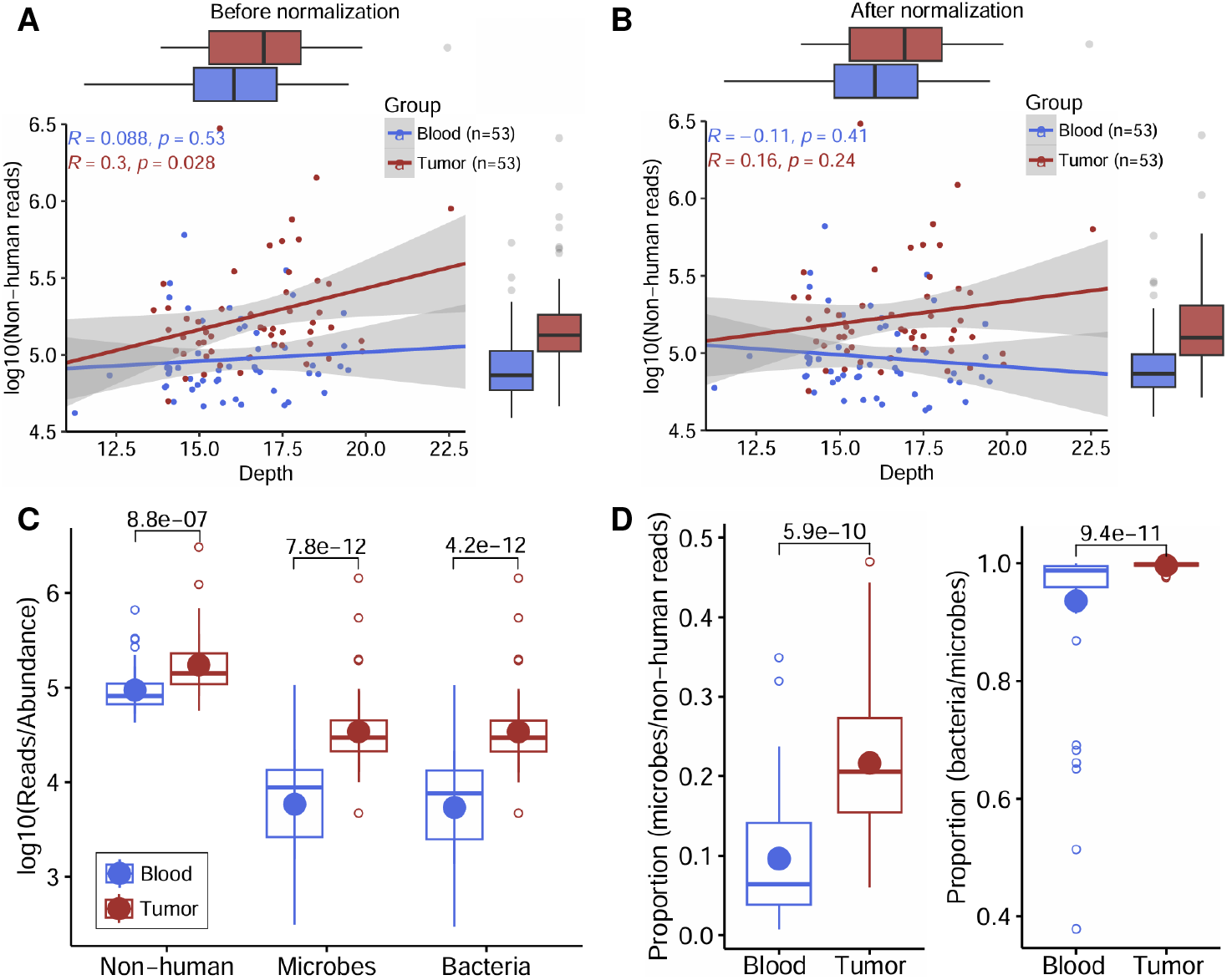
Comparisons of non-human reads, microbes, and bacteria in tumor and blood samples from DFSP patients. **(A, B)** Comparisons of non-human reads between tumor and blood samples before (**A**) and after **(B)**normalization. The points in red and in blue represent the tumor and blood samples, respectively. The lines in red and in blue refer to the fitted lines in tumor and blood, respectively. Spearman’s correlations between non-human reads and sequencing depths of human genome in tumor and blood were calculated, respectively. **(C)** Comparisons between tumor and blood across the levels of non-human reads, microbes, and bacteria. The line and point in each boxplot represent the median and mean, respectively. Significance was determined by two-tailed Wilcoxon test. **(D)** Comparisons of the proportions of microbes to non-human reads (**left**) and the proportions of bacteria to microbes (**right**) between tumor and blood. The line and point in each boxplot represent the median and mean, respectively. Significance was determined by two-tailed Wilcoxon test.

Further to determine which microorganisms the high-quality non-human reads belong to, we employed Kraken2 software in combination with PlusPF database for taxonomic annotation. ^[18, 19]^ To minimize false-positive microbial assignments originating from human sequences, the PlusPF database integrated human genome with a comprehensive microbial genome repository encompassing bacterial, viral, archaeal, and fungal kingdoms. ^[24]^ Here, the four kingdoms of bacteria, viruses, archaea, and fungi were considered as microbes in the tumor or blood samples. We consistently found that the abundance of non-human reads, microbes, and bacteria in tumor were significantly higher than those in blood (**Fig. 1C**). In particular, the proportions of microbes to non-human reads (**Fig. 1D**, left) and the proportions of bacteria to microbes (**Fig. 1D**, right) were also higher in tumor than those in blood. Previously, due to injury, infection, and barrier loosening, the microbiota in blood mainly originated from the translocation of those microbiota enriched in human body sites, such as oral, gut, and skin. ^[25]^ Therefore, we considered that those resident microbes in skin should exhibit higher detection rates than those in blood, indicating that those microbes in blood could be used to screen intratumor microbes (**Methods**; **Fig. S1A**). ^[23]^ Meanwhile, we observed that the median proportions of bacteria to microbes both showed over 98% (**Fig. 1D**, right), suggesting that the vast majority of microbes was consisted of bacteria in the tumor and blood samples. Thus, we only focused on bacteria in subsequent investigations.

Here, a total of 8,103 and 6,838 bacterial species were annotated in tumor and blood samples, respectively. As shown in **Fig. S1**, in order to identify key intratumor species in DFSP and minimize the impact of contaminants and noise, we referred to two prior studies and conducted a series of filtering procedures and treated blood samples as negative controls (**Methods**). ^[10,23]^ Consequently, in total, 916 intratumoral bacterial species were retained in the 53 tumors of DFSP (**Table S2**).

### Characteristics of intratumoral microbiome and their associations with malignancy in DFSP

Based on the abundance of these 916 intratumoral species identified in 53 tumor samples, we first evaluated their alpha-diversities, including Shannon index and species number (SpeciesNo). We found that Shannon (**Fig. 2A**) and SpeciesNo (**Fig. 2B**) were both negatively correlated with both Ki67 expression and tumor volume, indicating that the increased intratumoral microbial diversity may correspond to the decreased malignant in DFSP. Additionally, we estimated the Bray–Curtis dissimilarity between tumor samples. Then, principal coordinate analysis (PCoA) were performed to visualize beta-diversity, illustrating that those tumor samples with high alpha-diversity indexes appeared to be clustered (**Fig. 2C**).

**Fig. 2.**
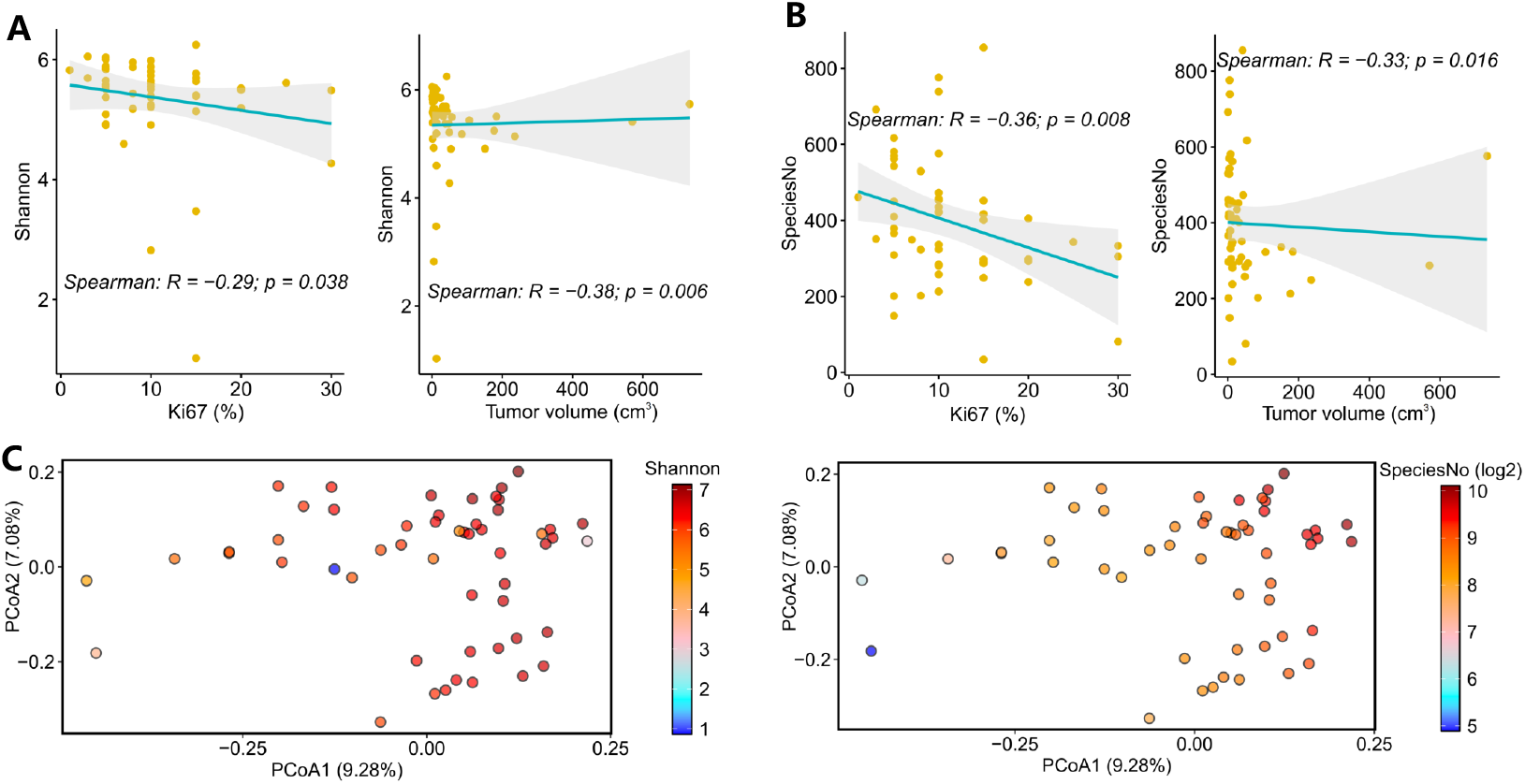
Alpha-diversity and beta-diversity of intratumoral species in DFSP. **(A)** Spearman’s correlations between Shannon and Ki67 expression (**left**) and tumor volume (**right**). Each point represents a tumor samples, and the line refer to the fitted line. **(B)** Spearman’s correlations between species number (SpeciesNo) and Ki67 expression (**left**) and tumor volume (**right**). Each point represents a tumor samples, and the line refer to the fitted line. **(C)** The beta-diversity based on the Bray–Curtis dissimilarity between samples. Each point represents a tumor sample, whose color is reflected by Shannon (**left**) or SpeciesNo (**right**), respectively.

These 916 intratumoral species distributed across 13 distinct phyla, 143 families, and 379 genera (**Table S3A-C**). At the phylum level, *Actinomycetota* and *Pseudomonadota* were found to be predominant in each sample (**Fig. 3A, top**), and both phyla exhibited negative correlations with Ki67 expression in DFSP (**Fig. 3B**). Furthermore, the top 10 families collectively accounted for a dominant proportion of microbial community in all samples (**Fig. 3A, middle)**, in which several, i.e. *Erythrobacteraceae, Streptomycetaceae, Xanthomonadaceae, Sphingomonadaceae, Micromonosporaceae, Pseudonocardiaceae*, and *Comamonadaceae*, demonstrated negative associations with Ki67 expression (**Fig. 3C**). Similarly, among the top 25 genera that represented the majority of microbial community **(Fig. 3A, bottom)**, we found that *Qipengyuania, Rhodoferax, Pseudonocardia, Pannonibacter*, and *Streptomyces* were negatively correlated with the expression of Ki67 (**Fig. 3D**). These findings supported a hypothesis that these dominant intratumoral bacteria may exert an inhibitory effect on the tumor malignancy, as indicated by the expression of Ki67 in DFSP.

**Fig. 3.**
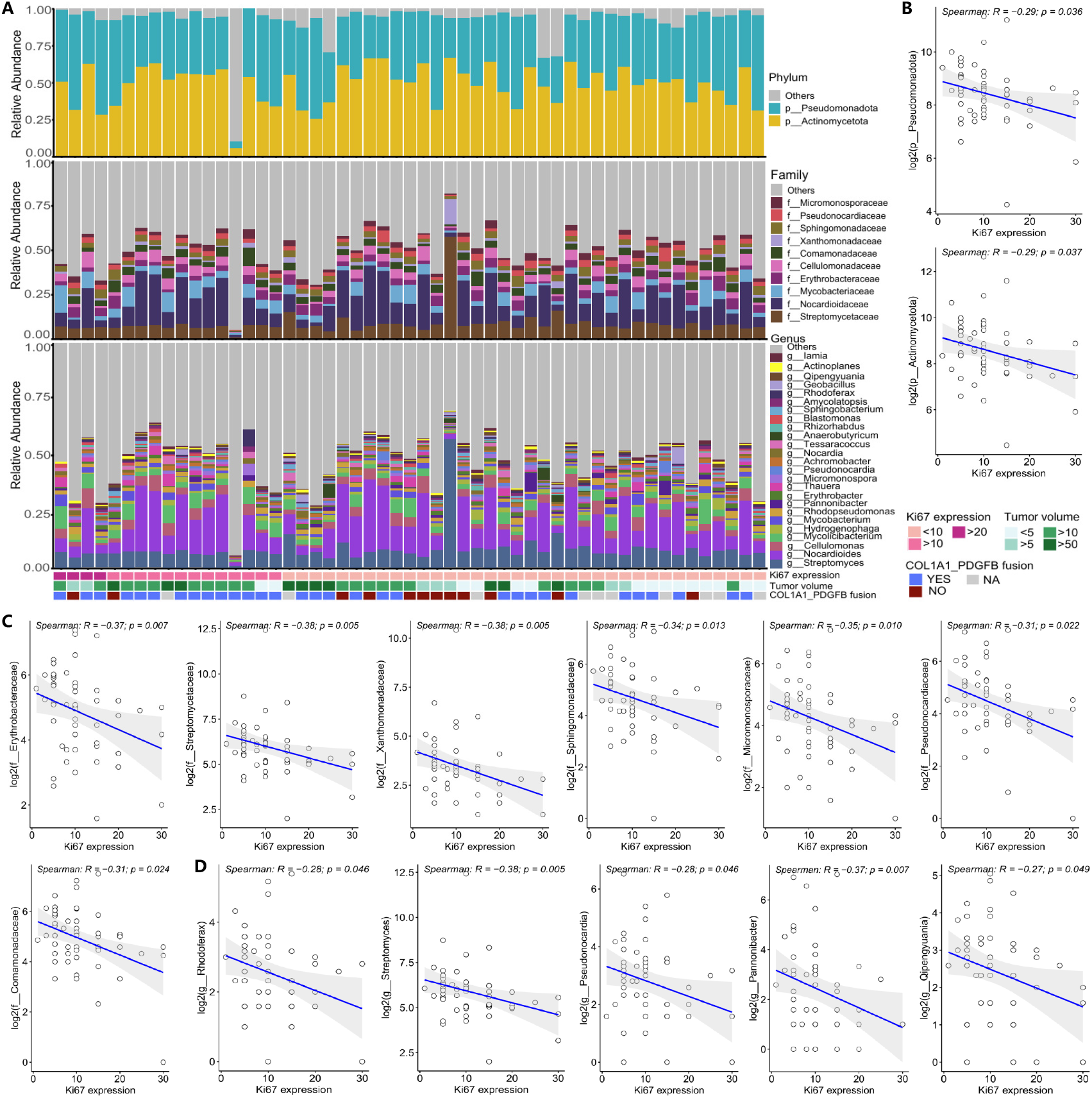
The predominant microbial taxa in DFSP and their correlations with Ki67 expression. **(A)** The distribution of the predominant microbial taxa across the phylum (**top**), family (**middle**) and genus levels (**bottom**). **(D)** Spearman’s correlations between Ki67 expression and the two predominant phyla. Each point represents a tumor samples, and the line refer to the fitted line. **(E)** Spearman’s correlations between Ki67expression and the five predominant families. Each point represents a tumor samples, and the line refer to the fitted line. **(F)** Spearman’s correlations between Ki67 expression and the five predominant genera. Each point represents a tumor samples, and the line refer to the fitted line.

### Identification of crucial intratumoral species associated with tumor malignancy of DFSP

Further to identify those crucial species in DFSP, those 916 intratumoral species were subjected to weighted gene co-expression network analysis (WGCNA). we evaluated a range of soft-threshold powers to construct scale-free topology network, and ultimately selected a power of 6 **(Fig. S2A)**. We verified that a scale independence (R^2^) of 0.92 and a connectivity slope of -1.49 were yielded, thereby confirming the network’s suitability for robust module detection **(Fig. S2B)**. Subsequently, we observed that all modules exhibited negative correlations with Ki67 expression and/or tumor volume **(Fig. S2C)**. Therein, the modules in yellow, in brown, in green, in blue, and in red showed statistically significant associations, and thus those intratumoral bacteria in these modules were selected.

Additionally, to identify the crucial species among 916 intratumoral bacteria, we again conducted gradient series analysis based on the distinct volume of tumor samples. As shown in **Fig. S2D**, we found that *Clusters 2, Cluster 15* and *Cluster 20* showed a marked decreasing trend with increasing tumor volume, thereby warranting the selection of intratumoral species within these clusters. Eventually, we intersected those species lists selected by the WGCNA and gradient series analyses above, and obtained 84 crucial species for subsequent analysis **(Fig. S2E)**.

As expected, most of these 84 species were observed to negatively correlate with tumor volume and/or Ki67 expression, e.g. *Streptomyces sp. P3, Streptomyces rutgersensis, Kutzneria albida, Sorangium cellulosum* and *Phreatobacter aquaticus* (**Fig. 4A**). Meanwhile, eight species were found to exhibit lower abundance in those samples with *COL1A1-PDGFB* fusion compared to those without, such as *Streptomyces sp. NEAU. sy36, Streptomyces sp. ITFR*.*21, Streptomyces rutgersensis, Caldimonas brevitalea, Hydrogenophaga taeniospiralis, Sphingorhabdus sp. YGSMI21, Altererythrobacter sp. B11, Sinirhodobacter sp. HNIBRBA609* (**Fig. 4B**). Moreover, nine species of them were found to exhibit lower abundance in those samples with trauma history compared to those without, such as *Streptomyces rutgersensis, Streptomyces sp. NEAU*.*sy36, Saccharothrix sp. 6*.*C, Pelagibacterium halotolerans, Pelagibacterium sp. 26DY04, Pseudolabrys taiwanensis, Pontivivens ytuae, Erythrobacter aurantius, Sphingosinithalassobacter sp. CS137, Opitutus sp. GAS368* (**Fig. 4C**). These indicated that several intratumoral species may have significant negative associations with the malignancy in DFSP.

**Fig. 4.**
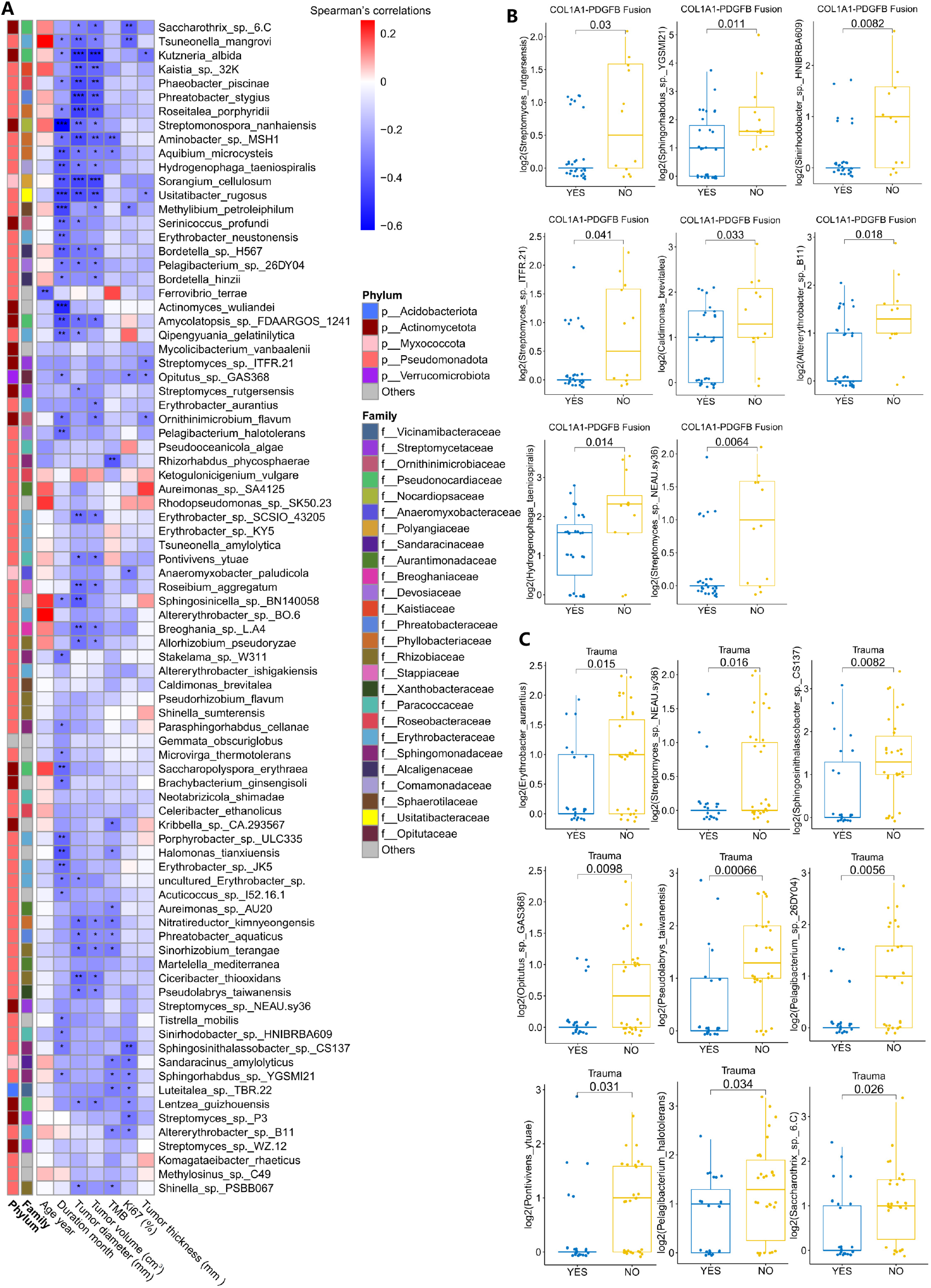
Correlation analysis between intratumoral species and clinical indicators in DFSP. **(A)** Heatmap illustrates the Spearman’s correlations between 84 crucial intratumoral species and several available clinical indicators, in which the cells in red or blue denote positive or negative correlations, respectively. * *P* < 0.05, ** *P* <0.01, *** *P* < 0.001. **(B)** Those significantly differential intratumoral species in those tumor samples with or without *COL1A1-PDGFB* fusion event. Significance was determined by two-tailed Wilcoxon test. **(C)** Those significantly differential intratumoral species in those tumor samples with or without a history of trauma. Significance was determined by two-tailed Wilcoxon test.

Furthermore, we also observed that some intratumoral bacteria exhibited significant negative correlations with TMB (**Fig. 4A**). Subsequently, further to investigate the impact of intratumoral bacteria on mutational types, we analyzed their correlations. As shown in **Fig. S3**, the general negative correlations were observed. Notably, we found that the predominant mutation type was single-nucleotide variation, with the dominant occurrence of T>C and T>G. That may echo to the observation of the negative correlations between intratumoral microbes and tumor malignancy in DFSP.

### Characteristics of crucial intratumoral microbiome associated with tumor malignancy of DFSP

To validate the above associations between those crucial intratumoral species and tumor malignancy of DFSP, we further clustered these 53 tumor samples based on the abundance of the 84 crucial intratumoral species. As shown in **Fig. 5A**, two distinct groups were formed owing to their difference in intratumoral species: *Group 1* with lower microbial abundance and *Group 2* with higher microbial abundance. By comparison, we revealed that those samples in *Group 2* exhibited significantly reduced Ki67 expression and smaller tumor volume than those in *Group 1* (**Fig. 5B**). And, a lower percentage of *COL1A1-PDGFB* fusion events was observed in *Group 2* compared to *Group 1* (**Fig. 5C**). Thus, these findings confirmed that the abundance of these crucial intratumoral species could inhibit the tumor malignancy of DFSP.

**Fig. 5.**
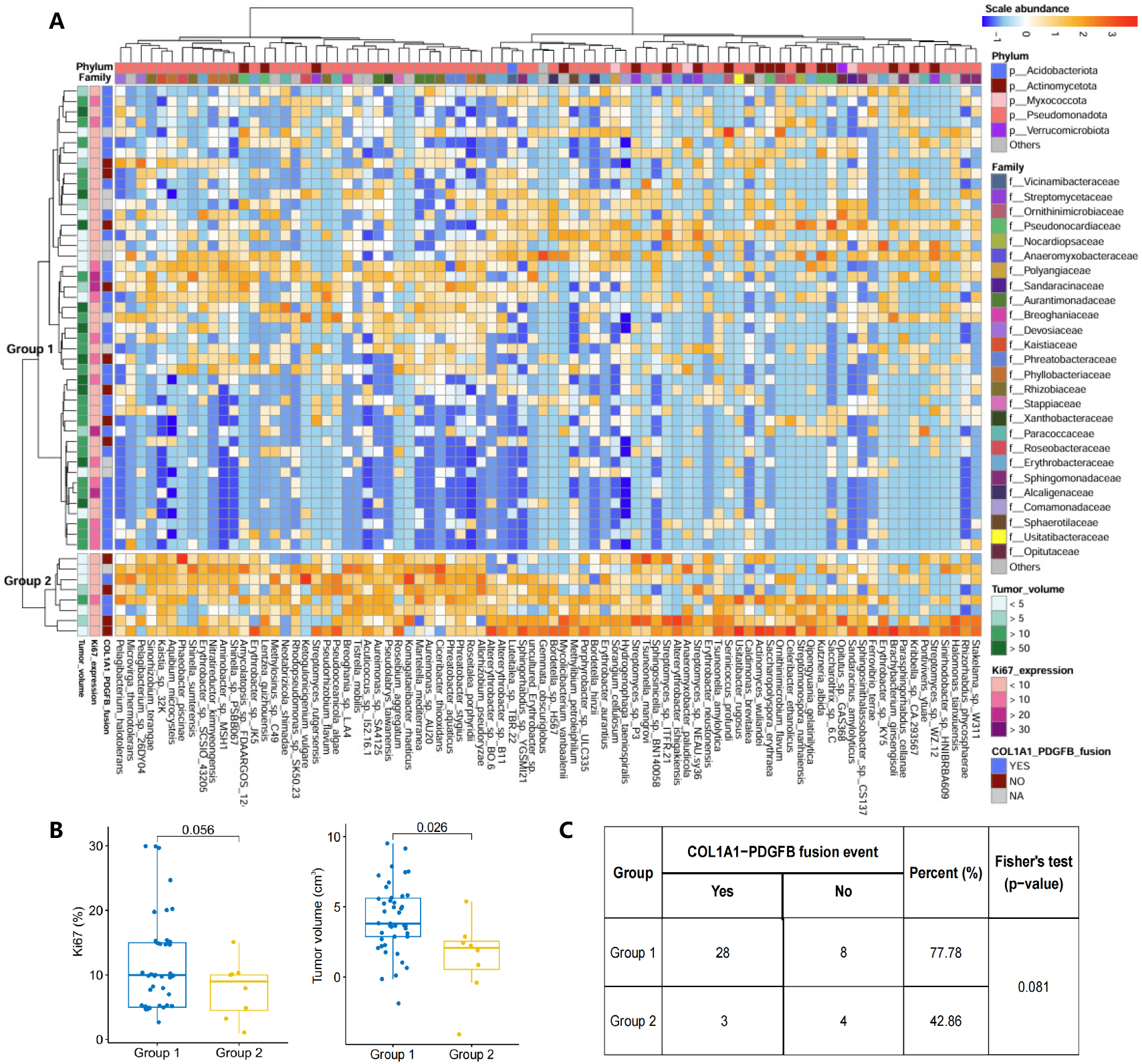
Clustering and comparisons of those tumor samples with high or low intratumoral species in DFSP. **(A)** Cluster analysis based on the abundance of the 84 crucial intratumoral species, in which the 53 samples were classified into two groups (i.e. *Group 1* and *Group 2*). Higher abundance was observed in *Group 2* compared to in *Group 1*. **(B)** Comparisons of Ki67 expression (**left**) and tumor volume (**right**) between *Group 1* and *Group 2*. Significance was determined by two-tailed t-test. **(C)** Comparison of *COL1A1-PDGFB* fusion event in *Group 1* and *Group 2*. Significance was determined by Fisher’s test.

## DISCUSSION

This study investigated, for the first time, the intratumoral microbiome in DFSP through analyzing WGS data from the paired biopsy and blood samples of DFSP patients. Microbial sequences were detected extensively in tumors and blood of DFSP patients. The levels of non-human reads, microbes, and bacteria were all significantly higher in tumor than those in blood, with the majority of microbes belonging to bacteria in both tumor and blood samples. We revealed the intratumoral microbiome in DFSP after filtering out contaminants and noise species, and found that the alpha-diversity was negatively associated with the malignancy of DFSP. Furthermore, the predominant intratumoral bacterial taxa were observed to exhibit negative correlations with Ki67 expression across the phylum (e.g. *Actinomycetota*), family (e.g. *Streptomycetaceae*) and genus (e.g. *Streptomyces*) levels. And, several crucial species (e.g. *Streptomyces rutgersensis, Streptomyces sp. P3*, and *Kutzneria albida*) were found to significantly correlate with the diminished Ki67 expression, reduced tumor volume, and decreased *COL1A1-PDGFB* fusion events. Together, these findings suggested the potential suppressive roles of the dominant intratumoral bacteria in the malignancy of DFSP.

Indeed, *Kutzneria albida* has emerged as a highly promising microbial strain for the synthesis of vitamin D2 and vitamin D3. ^[26]^ These vitamins are pivotal in regulating cellular processes within the human body and are potentially effective in the treatment of several diseases, including autoimmune diseases and cancers. ^[27-30]^ *Streptomyces* species have long been recognized for their anti-tumor properties, with prior research linking their enrichment in pancreatic cancer to prolonged survival via enhanced anti-tumor immunity and metabolic reprogramming. ^[31]^ Members of *Streptomycetaceae* family were considered as prolific antibiotic producers, with compounds such as streptonigrin exhibiting broad-spectrum anticancer activity and neoantimycin F (NAT-F) inducing apoptosis in NSCLC cells. ^[32-37]^ That is to say, we hypothesized that these dominant intratumoral bacteria in DFSP may suppress tumor malignancy by modulating the immune system and/or interfering tumor cell metabolism.

Additionally, various somatic mutations, especially the translocation between chromosomes 17 and 22, are crucial for the tumorigenesis of DFSP. ^[38, 39]^ Herein, we observed the negative correlations between intratumoral species and single-nucleotide variations (e.g. T > C, T > G), which is the predominant mutation type in DFSP. That suggested that intratumoral bacteria in skin may promoting immune recognition and clearance of mutational cells, thereby contributing to the inhibition of malignancy in DFSP.

Our study has several limitations. Firstly, the WGS data of paired tumor and blood samples from DFSP patients was simultaneously used to investigate intratumoral microbiome associated with DFSP, but this study was limited by the use of a single dataset and required to be validated by other independent datasets. Furthermore, although we proposed and conducted a series of filtering of contaminants and noise bacteria, there remains the possibility of false positives being retained in those intratumoral species. Lastly, the clinical significance of our findings and the underlying mechanisms of intratumoral species in DFSP still required further experimental validation to confirm their implications.

Overall, this study performed, for the first time, a comprehensive investigation of the intratumoral microbiome in DFSP, in which we unveiled the pivotal role of intratumoral microbes in mitigating tumor malignancy and provided a novel insight for DFSP pathogenesis and clinical management.

## Acknowledgments

We appreciate the resources provided by the High Performance Computing Center of Central South University.

## Contributors

XJ conceived and designed the study. ZL, JZ, and YL contributed to investigation and methodology. CP, MW, and XC are dermatologist and experts, who provided professional guidance and sequencing data. GZ, BL, JL, and LX provided the high-performance computing platform and bioinformatics guidance. XJ and ZL carried out data analysis, validation and visualization. ZL wrote the original draft and XJ edited this manuscript under the supervision of CP, MW, and XC. XJ conceived the project and provided funding. All authors approved this final version for the decision to submit for publication.

## Data availability

No original sequences were produced in this study. The WGS datasets analyzed herein were derived from a previously published study. ^[17]^ The outcomes of microbial annotation generated in this study were stored in a public database National Omics Data Encyclopedia (NODE), with access number OEZ00020956 (https://www.biosino.org/node/review/detail/OEV00000624?code=CA4SMTKM).

## Ethics declarations

The study was approved by the ethics committees of Xiangya Hospital of Central South University, and the written informed consent from all participants or their legal guardians were obtained.

## Consent for publication

Not applicable.

